# Genome diversification of symbiotic fungi in beetle-fungus mutualistic symbiosis

**DOI:** 10.1101/2024.10.09.617500

**Authors:** Yin-Tse Huang, Khaled Abdrabo El-Sayid Abdrabo, Guan Jie Phang, Yu-Hsuan Fan, Yu-Ting Wu, Jie-Hao Ou, Jiri Hulcr

## Abstract

Ambrosia beetles and their fungal symbionts represent a widespread and diverse insect-fungus mutualism. This study investigates the genomic adaptations associated with the evolution of the ambrosia lifestyle across multiple fungal lineages. We performed comparative genomic analyses on 70 fungal genomes from four families (Irpicaceae, Ceratocystidaceae, Nectriaceae, and Ophiostomataceae), including 24 ambrosia and 34 non-ambrosia lineages. Our phylogenomic analyses reveal multiple independent colonization of insect vectors by the fungi, spanning from the mid-Cretaceous (114.6 Ma) to the early Quaternary (1.9 Ma). Contrary to expectations for obligate symbionts, ambrosia fungi showed no significant genome-wide reductions in size, gene count, or secreted protein repertoire compared to their non-symbiotic relatives. Instead, we observed conservation of most assessed genomic features; where genome traits differ between free-living relatives and ambrosia fungi, the changes are lineage-specific, not convergent. Key findings include lineage-specific expansions in carbohydrate-active enzyme families (AA4 in Nectriaceae, CE4 in Ophiostomataceae, and GH3 in Ophiostomataceae and Ceratocystidaceae), suggesting potential enhancement or loss of lignin modification, hemicellulose deacetylation, and cellulose degradation in different ambrosia lineages. Repeat-Induced Point mutation analysis revealed family-specific patterns rather than lifestyle-associated differences. These results highlight the diverse genomic strategies employed by ambrosia fungi, demonstrating that symbiont evolution can proceed through refined, lineage-specific changes rather than genome-wide, or convergent alterations. Unlike other insect-associated fungi, ambrosia fungi do not seem to be domesticated crops, but rather free-living fungi which co-opted wood boring beetles as vectors via subtle, lineage-specific adaptations.

## Introduction

Insect-fungal symbioses include remarkably stable and successful evolutionary strategies with ecological significance. Insects and fungi engage in interactions ranging from mutualistic to commensal to competitive, and are pivotal in nutrient cycling and decomposition of wood. Among these relationships, the ambrosia symbiosis is the most widespread, diverse, and repeatedly evolved. It involves the wood-boring ambrosia beetles in the weevil subfamilies Scolytinae and Platypodinae, and their fungal partners from several fungal orders. Ambrosia beetles vector fungi between dead trees - the fungal substrate - and feed on fungal conidia within the tunnels. The fungi benefit from beetle-assisted transport to new tree hosts. This mutualism shows co-evolution, with beetles evolving mycangia to carry fungal tissue ^1,2^, and fungi adapting ambrosial cells to for nutrient provisioning to the beetles ^3–5^. Both partners have specialized adaptations supporting their interdependence.

Both the ambrosia beetles and the ambrosia fungi exemplify convergent evolution, with multiple independent origins across different lineages ^2,6,7^. The mutual evolved at least 12-16 times in the scolytine and platypodine beetles ^7,8^, and similarly over 10 times in both Basidiomycota and Ascomycota fungi ^9,10^. These multiple instances of parallel evolution provide an opportunity to study genomic adaptations supporting mutualistic interactions across independent evolutionary backgrounds.

Recent genomic studies on various fungal symbionts outside of the beetle system, including mycorrhizal fungi and fungal crops of insects, have revealed insights into genome evolution and functional gene diversification ^11–14^. However, most such symbioses evolved only once. Ambrosia fungi and their multiple evolutionary origins offer unparalleled designs for comparative phylogenetic analyses.

Despite the appearance of uniformity of the ambrosia lifestyle, recent studies show that ambrosia fungi’s metabolomic profiles don’t converge across lineages ^3,15^. This lack of metabolomic convergence, combined with a strong influence of ancestral metabolic features, suggest the uniqueness of each ambrosial system, and beckons redefining what ambrosia fungi are. Ambrosia fungi are not internal symbionts, restricted to an environment controlled by their animal partner. Instead, they spend most of their life cycle outside of the animal symbiont, occupying their ancestral niche in freshly dead trees. Thus, compared to the genomes of their free-living relatives, the genomes of ambrosia fungi are unlikely to evolve major differences in metabolism, catabolism, or secondary compounds. Just as their relatives, ambrosia fungi can use wood polymers as carbon sources ^16^ but most are non-competitive degraders of lignocellulose compared to true wood decay fungi (Skelton et al., 2019).

The two key fungal adaptations to the ambrosia lifestyle, as far as is known, include the colonization of the beetle’s transport receptacle, the mycangium, and the capacity to sequester nutrients necessary for the beetle development. There is currently no robust theory predicting what metabolic pathway would be up-or down-regulated to achieve these adaptations, or any other traits facilitating the symbiosis. The adaptations might be subtle ^17^. They may also be varied across different fungal lineages due to their multiple independent origins and subsequent specificity to their beetle lineages (Hulcr and Stelinski, 2017).

Functional genes, particularly those encoding carbohydrate-active enzymes (CAZymes), lipases, peptidases, and cell wall-degrading enzymes, are crucial indicators of fungi’s metabolic capabilities and ecological roles ^18,19^. However, recent research on ambrosia fungi suggests that their enzymatic profiles may be adapted differently than previously thought.

Rather than being efficient at plant biomass degradation, these fungi appear to be specialized for rapid utilization of simple nutrients available in their beetle galleries ^17,20^. Therefore, key questions remain about the nature and direction of specialization in the enzymatic profiles of ambrosia fungi, and how these profiles compare to those of their non-symbiotic relatives.

One of the important features of fungal molecular ecology are G-protein coupled receptors (GPCRs), crucial for fungi to sense and respond to environmental cues, detecting signals such as nutrients, pheromones, and stress factors. These transmembrane proteins regulate fungal development, metabolism, and pathogenicity ^21,22^. In pathogenic fungi, certain GPCR families, like Pth11-like receptors, are key for host recognition and infection ^23,24^. However, the role and evolution of GPCRs in symbiotic fungi, particularly in ambrosia symbiosis, remain unexplored. Understanding the GPCR repertoire in ambrosia fungi could reveal how these symbionts perceive and interact with their beetle hosts.

Another potentially critical component of ambrosia fungal genome are candidate secreted effector proteins. These specialized proteins are likely instrumental in mediating host interactions, particularly in the establishment and maintenance of the symbiotic relationship with their hosts ^25–27^. By analogy with other fungal symbioses, such as mycorrhizal associations, these effectors may facilitate nutrient exchange and modulate host immune responses, both of which may be important in the colonization of the beetle mycangium.

Repeat-Induced Point mutation (RIP) is another crucial aspect of fungal genome evolution that warrants investigation in ambrosia fungi. This genome defense mechanism, which mutates and silences repetitive DNA sequences, can significantly influence genome architecture and evolution. While RIP is typically associated with sexual reproduction, ambrosia fungi are thought to be primarily asexual ^28–31^. However, sporadic discoveries of sexual states in some ambrosia lineages ^32,33^ suggest the potential for RIP activity through cryptic sex.

Our study explores the genomic foundations underlying ambrosia fungi’s adaptation to their specialized ecological niche. We compared 70 fungal genomes across four families (Irpicaceae, Ceratocystidaceae, Nectriaceae, and Ophiostomataceae), including 24 ambrosia and 34 non-ambrosia lineages, plus 12 outgroups. This dataset, comprising 8 newly sequenced and 62 retrieved genomes, represents independent origins of ambrosia fungi. Our research aims to uncover distinctive genomic characteristics associated with the ambrosia lifestyle by addressing five key questions: 1) How do the genomic features of ambrosia fungi compare to those of other fungal symbionts and their non-symbiotic relatives? 2) What specific enzymatic adaptations correlate with the origins of the symbiotic relationship with ambrosia beetles? 3) How do secreted proteins and GPCRs contribute to mutualistic recognition in the ambrosia symbiosis? 4) What role does Repeat-Induced Point mutation (RIP) play in shaping the genome architecture of ambrosia fungi? This study explores the evolutionary patterns, genomic foundations, and ecological adaptations of ambrosia fungi, advancing our understanding of insect-fungal symbiosis from mere observations to genome-level understanding.

## Results and Discussion

### Genome Assembly and Quality Assessment

We sequenced genomes of eight species in the Ophiostomataceae, Ceratocystidaceae, Nectriaceae, and Irpicaceae families, including both ambrosia and non-ambrosia fungi, and obtained genomes of 62 additional fungal species from the NCBI Genome database. Genomic features are summarized in Table S2. All genomes showed Benchmarking Universal Single-Copy Orthologs (BUSCO) values above 96%, indicating high-quality assemblies (Figure 1). Genome sizes varied across families, ranging from 28 Mb (*Ambrosiella roeperi* T.C. Harr. & McNew) to 54.4 Mb [*Neocosmospora euwallaceae* (S. Freeman, Z. Mendel, T. Aoki & O’Donnell) Sand.-Den., L. Lombard & Crous)]. GC content was relatively consistent [45% in A. roeperi to 55.3% in *Harringtonia lauricola* (T.C. Harr., Fraedrich & Aghayeva) Z.W. de Beer & M. Procter)]. Assembly contiguity varied considerably, with N50 values ranging from 0.38 Mb [(*Irpex subulatus* (Ryvarden) Z.B. Liu & Y.C. Dai)] to 4.45 Mb (*N. euwallaceae*). Scaffold counts ranged from 39 [(*Leptographium procerum* (W.B. Kendr.) M.J. Wingf., Trans)] to 476 (*I. subulatus*). Protein-coding gene counts showed substantial variation, from 8,158 [(*Huntiella moniliformis* (Hedgc.) Z.W. de Beer, T.A. Duong & M.J. Wingf.)] to 21,224 (*N. euwallaceae*). Notably, Nectriaceae family members [(*N. euwallaceae* and *N. solani* (Mart.) L. Lombard & Crous)] exhibited higher gene counts compared to other families.

**Figure 1.**
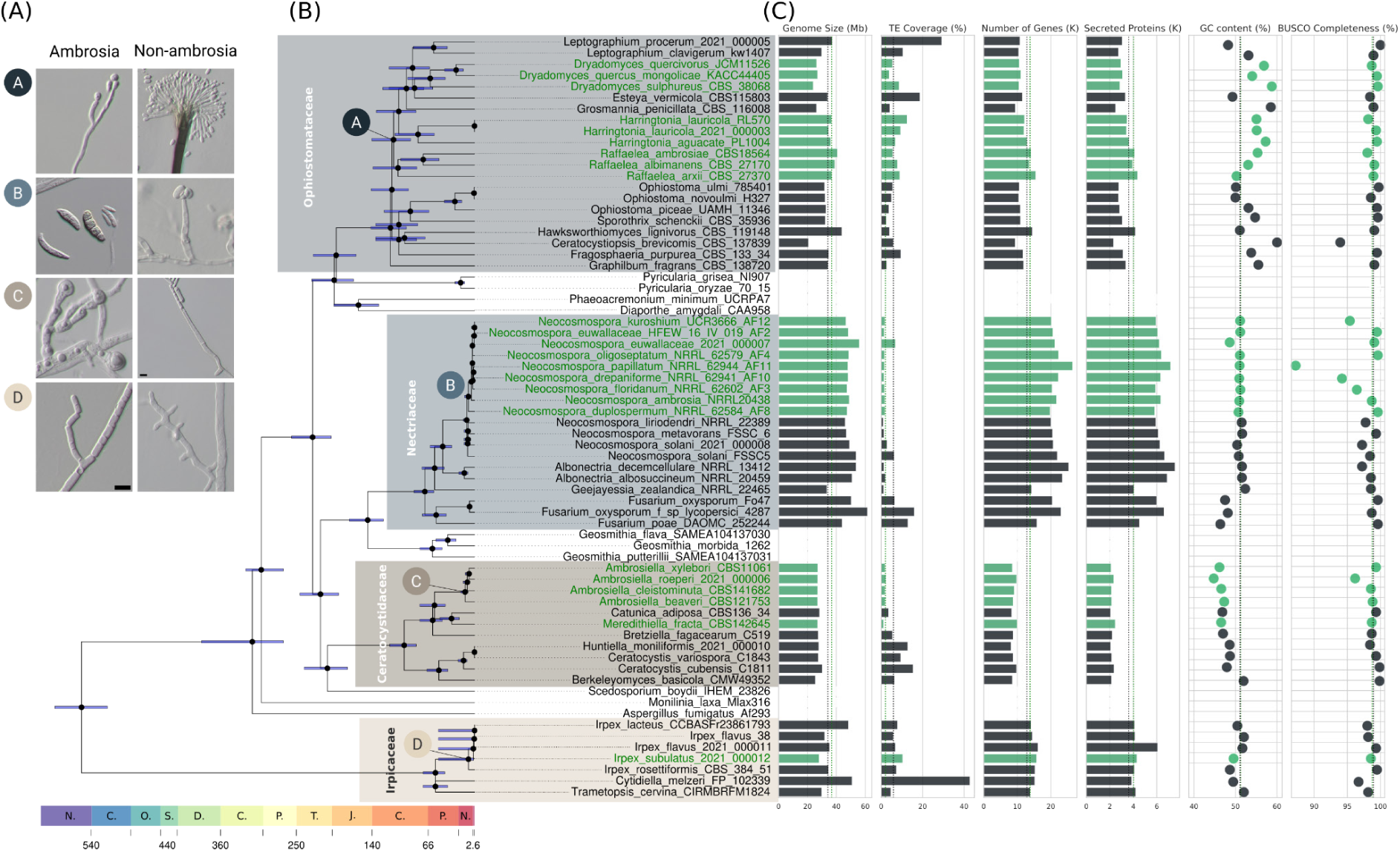
Evolution and genomic features of ambrosia fungi. (A) Representative fungal species illustrate morphological adaptations (enlarged cells) in ambrosia vs. non-ambrosia fungi; colored circles denote representative species in four fungal families. (B) Dated phylogenomic tree showing multiple origins of ambrosia fungi across Ascomycota and Basidiomycota. Green taxa represent ambrosia fungi. Time scale in millions of years ago (Ma) shown at the bottom. (C) Comparison of key genomic features (genome size, TE coverage, gene count, and secreted protein count) between ambrosia (green) and non-ambrosia (black) fungi. Outgroup species not shown.

No consistent genomic patterns (Figure 2 & Table 2) were observed between ambrosia and non-ambrosia fungi across families, implying that the ambrosia lifestyle may not be associated with convergent genome-wide characteristics.

**Figure 2.**
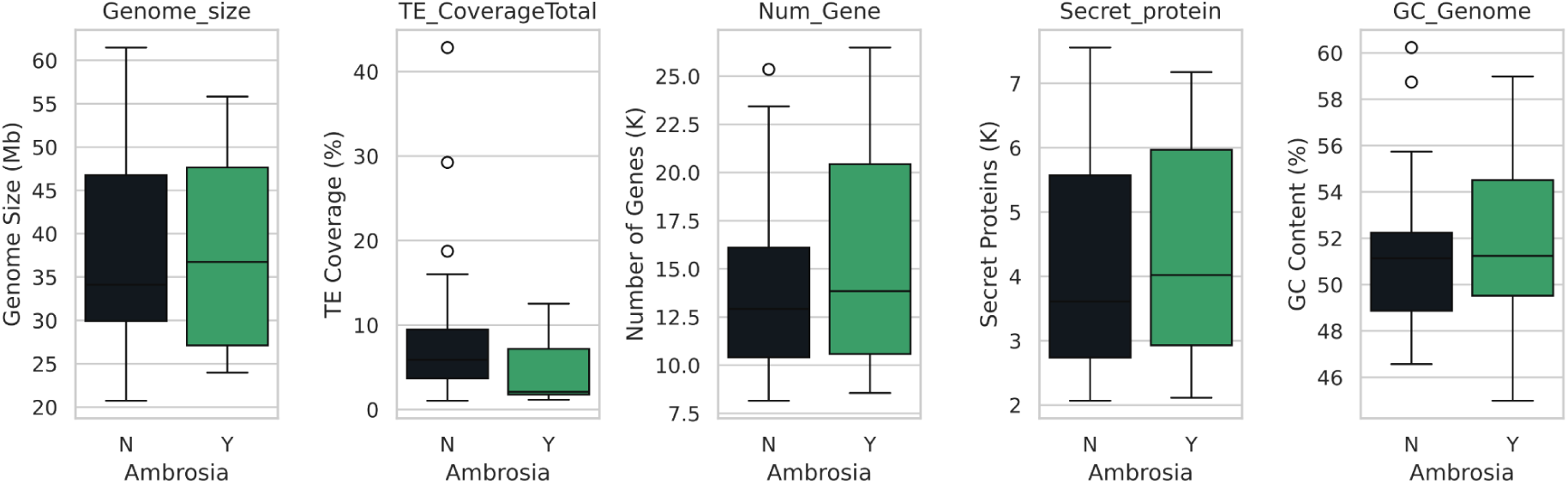
Comparison of genomic features between ambrosia (green) and non-ambrosia (black) fungi. No statistically significant differences were found for genome size, transposable element coverage, gene count, secreted protein count, or GC content across all species tested.

### Multiple symbiosis events of ambrosia fungi across Dikarya

Our phylogenomic analyses, based on genome-wide sequence data from 70 taxa—including 24 ambrosia fungi, 34 non-ambrosia relatives, two Mucoromycota outgroups (not shown in the time-calibrated phylogeny), and ten taxa from the Eurotiomycetes, Leotiomycetes, and Sordariomycetes groups—enhanced the resolution of the phylogenetic topology. The resulting phylogenomic tree indicates that ambrosia fungi colonized the insect vectors repeatedly, over a wide evolutionary period, spanning from the mid-Cretaceous (114.6 Ma) to the early Quaternary (1.9 Ma) (Table 1). This tree reveals multiple independent beetle colonization events and subsequent co-diversification (Figure 1 & S1).

**Table 1.** Divergence dates of ambrosia lineages in the present study.

| Ambrosia species | Family | Crown age (Ma) | CI_Lower (Ma) | CI_Upper (Ma) | Stem age (Ma) |
| --- | --- | --- | --- | --- | --- |
| *Dryadomyces* spp. | Ophiostomataceae | 66.41 | 44.07 | 100.06 | 102.03 |
| *Harringtonia* spp. | Ophiostomataceae | 84.69 | 60.21 | 119.12 | 113.46 |
| *Raffaelea* spp. | Ophiostomataceae | 114.62 | 86.44 | 152.00 | 121.31 |
| *Neocosmospora* spp. | Nectriaceae | 9.37 | 5.80 | 15.14 | 11.82 |
| *Ambrosiella* spp. | Ceratocystidaceae | 14.31 | 9.90 | 20.70 | 60.08 |
| *Meredithiella* spp. | Ceratocystidaceae | 34.60 | 22.98 | 52.10 | 60.08 |
| *Irpex subulatus* | Irpicaceae | 1.94 | 1.58 | 54.23 | 9.24 |

### Ancient Origins and Asynchronous Evolution in Ophiostomataceae

The phylogenetic topology of Ophiostomataceae in our study largely aligns with previous findings, albeit with some discrepancies. Notably, our placement of *Graphilbum fragrans* (Math.-Käärik) Z.W. de Beer, Seifert & M.J. Wingf. as the basal lineage within the Ophiostomataceae supports the findings of Nel et al. ^34^ and Vanderpool et al. ^35^. This differs from the topology presented in de Beer et al. ^36^, where *G. fragrans* is nested with *Leptographium*, *Dryadomyces*, and *Esteya* species. Consistent with earlier studies, our phylogeny indicates *Raffaelea* sensu stricto species clustered with a clade comprising *Leptographium*, *Dryadomyces*, *Harringtonia*, *Esteya*, and *Grosmannia* species, albeit with moderate support. While our analysis yielded 100% bootstrap support for this grouping, it is important to note that concatenated supermatrix datasets can sometimes artificially inflate support values, potentially masking underlying uncertainties ^37,38^. Gene and site concordance factors (gCF/sCF) for this node (6.68/34 respectively) closely mirror those reported by de Beer et al. ^36^ (10.1/34.6). Our incomplete lineage sorting (ILS) tests (Table S3) revealed significant values for both gCF (p = 3.85E-04) and sCF (p = 4.27E-07), suggesting that introgression or incomplete taxon sampling may have influenced this branching pattern. Future studies incorporating more closely-related species are needed to provide better resolution of their evolutionary relationships. Consistent with previous research, our phylogeny also shows a well-supported clade comprising *Ophiostoma*, *Sporothrix*, *Fragosphaeria*, and *Ceratocystiopsis*.

The Ophiostomataceae exhibit variable degrees of adaptation to beetles, characterized by high levels of promiscuity in beetle host associations (Li et al., 2018; Simmons et al., 2016). Our latest estimates suggest earlier divergence times compared to previous studies, pushing the origin of the first ambrosia symbioses deeper into the past.

The *Dryadomyces* clade (corresponding to the *Raffaelea sulphurea* complex or RS clade in Vanderpool et al., 2018) diverged approximately 66.4 million years ago (Ma) (95% CI: 44.1– 100.1 Ma), with a stem age of 102 Ma. This is considerably earlier than the 33 Ma crown age and 58 Ma stem age previously reported. The *Harringtonia* clade (equivalent to the complex previously named after *Raffaelea lauricola* or RL clade) shows an even older divergence at 84.7 Ma (95% CI: 60.2-119.1 Ma), surpassing the previous estimate of 67 Ma for its crown age. Notably, the *Raffaelea* sensu stricto clade (corresponding to the *Raffaelea ambrosiae* complex or RA clade) exhibits the earliest divergence at 114.6 Ma (95% CI: 86.4–152 Ma), with a stem age of 121.3 Ma. This substantially predates the 86 Ma crown age previously reported. It also aligns closely with estimates of the origin of Platypodinae (Coleoptera, Curculionidae), the oldest clade of ambrosia beetles, at ∼96 Ma by Jordal ^39^, but it predates the estimate by Shin et al. ^40^.

These deep divergence times, particularly in the *Raffaelea* clade, suggest that the lineages leading to these ambrosia fungi were already distinct around or before the estimated origin of fungus-feeding in some beetle groups. While beetle mycangia morphology has been suggested to influence symbiont fidelity ^41–43^, the frequent colonization of different ambrosia beetle lineages by Ophistomataceae may reflect aspects of fungal ecology rather than beetle morphology ^44^. Some fungal traits that are beneficial or conducive to transmission within the ambrosia symbiosis may have evolved prior to their association with beetles, predisposing these fungi to readily establish symbiotic relationships ^45^. This is consistent with the observation that different beetle lineages may have been independently colonized by various Ophiostomataceae fungi at different times, leading to the observed pattern of asynchrony. This revised evolutionary scenario provides a more nuanced understanding of the complex and ancient nature of these symbiotic relationships, emphasizing the importance of fungal pre-adaptations in directing the ambrosia symbiosis.

### Specialized Symbioses and Evolutionary Synchronization in Ceratocystidaceae

In contrast to the promiscuous nature of Ophiostomataceae, Ceratocystidaceae fungi exhibit higher fidelity to their beetle vectors. The *Ambrosiella*-*Meredithiella* clade diverged 34.6 Ma (95% HPD: 22.9–52.1 Ma), with crown ages closely approximating the origins of their respective hosts (Xyleborini and *Corthylus*) ^46^.

The Ceratocystidaceae is a promising model for comparative evolutionary analysis of the symbiosis evolution. Several genera of ambrosia fungi evolved independently in this clade, and all are highly specific to their beetle vectors (unlike many of the Ophistomataceae) ^33,43,47,48^. The fungus-beetle evolutionary units differ slightly in their relationships to host trees. Species within the *Ambrosiella*/*Xylosandrus* clade are polyphagous in terms of the tree host taxonomic identity, which contributes to their success as invasive species in many parts of the world ^49–51^. They are, however, highly specific to a particular stage of the tree tissue, death, and decay. This ecological niche—freshly dead or live but internally compromised tree branches—conforms to that of the necrotrophic phytopathogenic relatives of *Ambrosiella*, and it seems to have driven the adaptation of the beetle’s semiochemical ecology and host-selection behavior ^52^. The *Meredithiella* lineage, associated with various Corthylini beetles, shows similarly high mutual fidelity. However, *Meredithiella* fungi appear to be more specialized in terms of tree host range, as many species are found in specific hardwood tree species ^53^. Testing the extent to which these joint ecologies are driven by the fungal or the beetle ancestral ecology will require a more comprehensive sample of phylogenetic relatives of both the fungi and the beetles. Genome sequences of *Wolfgangiella*, *Phialophoropsis*, and *Toshionella* remain unavailable.

The contrasting host specificity between Ophiostomataceae and Ceratocystidaceae highlight the diversity of evolutionary strategies in ambrosia symbioses, with Ophiostomataceae retaining a more flexible, promiscuous relationship to both beetles and trees, while Ceratocystidaceae showing specialized ecologies.

### Adaptability and Promiscuity in Nectriaceae Ambrosia Fungi

In Nectriaceae, the *Neocosmospora* ambrosia clade (henceforth NAC, formerly known as the Ambrosia *Fusarium* Clade or AFC) presents a unique case of fungal adaptation to the ambrosia lifestyle. This monophyletic clade diverged approximately 9.4 Ma (95% HPD: 5.8– 15.1 Ma), aligning closely with the previous estimation (∼19.3 Ma; 95% HPD: 10.7–28.2 Ma) in O’Donnell et al. ^54^ and the divergence of their primary hosts, the *Euwallacea* beetles (∼ 12 Ma) ^46^. This near-synchronous divergence suggests a co-evolutionary relationship, yet not as strict as observed in some other ambrosia symbioses.

Like *Ambrosiella*, the NAC fungi are specific to the stage of tree death, but promiscuous in terms of tree taxonomy. The few studied species are parasites on stressed but living trees, or colonizers of moribund tissues, but rarely they colonize wood in advanced stages of death and decay ^54–57^. They are able to associate with multiple beetle vectors colonizing the same type of tree tissue. For instance, in Taiwan, NAC species such as AF-18 were found to be associated with three different *Euwallacea* species [Polyphagous Shot Hole Borer (PSHB), Tea Shot Hole Borer (TSHB), and Kuroshio Shot Hole Borer (KSHB)], while individual beetle species like PSHB were found to vector multiple NAC species (AF-13 to AF-18, and *N. kuroshium*) ^57^. Initial reports of strict associations in invaded areas, such as *N. euwallaceae* with *E. fornicatus* and *N. kuroshium* with *E. kuroshio*, may not represent the final specificity, but possibly the founder effect of limited initial populations.

The capacity to colonize partially living trees appears to be a native ecology of some, but not all, *Neocosmospora*/*Euwallacea* pairs ^58^. These species have become significant invasive pests worldwide, causing substantial damage to both native and cultivated trees ^59–61^. For example, *E. fornicatus* in California and Israel can attack over 200 woody plant species from 58 families, with about 32 species serving as reproductive hosts ^5^. The damage by these invasions may be partly due to the ability of their NAC symbionts to colonize and thrive in diverse tree species, although the tree death is attributable to the accumulation and persistence of attacks rather than fungal pathogenicity ^5^.

### Recent Divergence of Irpicaceae Ambrosia Fungi

A unique case is presented by *I. subulatus* (Irpicaceae, formerly known as *Flavodon subulatus* or *Flavodon ambrosius*), which diverged approximately 1.9 Ma (95% HPD: 1.6-54.2 Ma) according to phylogenomic analysis. This divergence time significantly postdates the estimated origin of its associated beetle hosts in the genera *Ambrosiodmus* and *Ambrosiophilus*, which evolved around 19 Ma ^46^. This temporal discrepancy suggests an asynchronous adaptation of the beetle and the fungus, potentially involving beetle lineages feeding on free living wood-decay fungi, later followed by a delayed evolution of the fungal adaptations to dispersal via the insect vector. It may also be a due to an undersampled phylogeny, reflecting the fact that basidiomycete ambrosia fungi are poorly sampled and virtually unstudied, even though some are associated with the most basal clades of scolytine beetles ^62^.

Notably, *I. subulatus* can produce free-living fruiting bodies ^63^, a trait analogous to *Termitomyces* fungi in termite-fungus farming systems ^64^. This capacity for environmental existence outside of the insect gallery, also observed in other ambrosia fungi like *R. lauricola* ^65^, may confer ecological advantages, allowing *I. subulatus* to survive without its beetle hosts and colonize new niches, akin to the situation in *Amylostereum*, a basidiomycete in which some populations are dispersed vertically by symbiotic wood-wasp ^66^, and some are dispersed horizontally ^67^. Such trait retention potentially influences the fungus’s population genetics, dispersal capabilities, and evolutionary trajectory, contributing to its success across multiple beetle host species and geographic regions ^68^. The apparent evolutionary mismatch between *I. subulatus* and its hosts presents an intriguing scenario for studying the dynamics of symbiont acquisition and co-evolution in ambrosia systems, particularly in cases where the symbiont may have been acquired more recently in evolutionary time. Additional ecological, biological, and genomic data on *Irpex* ambrosia fungi from their native range would greatly enhance our understanding of their evolutionary trajectories.

### Genome Features of Ambrosia Fungi Show Family-Specific Differences Without Broad Phylogenetic Trends

Our Phylogenetically Independent Contrasts (PIC) analysis revealed no statistically significant associations between the ambrosia lifestyle and the examined genomic features when accounting for phylogenetic relatedness (Figure 2 & Table 2). The analysis included genome size, transposable element (TE) coverage, number of genes, number of secreted proteins, and GC content. None of these features showed a significant difference between ambrosia and non-ambrosia fungi (Table 2, all p-values > 0.05). These results suggest that, contrary to expectations derived from studies of other symbiotic microorganisms, there is no clear genomic convergence associated with the ambrosia lifestyle across different fungal lineages. Adaptations to the ambrosial lifestyle might be lineage-specific rather than following a general pattern. This aligns with observations in other symbiotic systems, where genomic changes often reflect the unique evolutionary trajectories and ecological contexts of specific lineages ^69–71^. In ambrosia fungi, as in other symbiotic systems, these adaptations may be shaped by factors such as the host tree species, competing microorganisms, and specific environmental conditions ^72–74^.

**Table 2.**
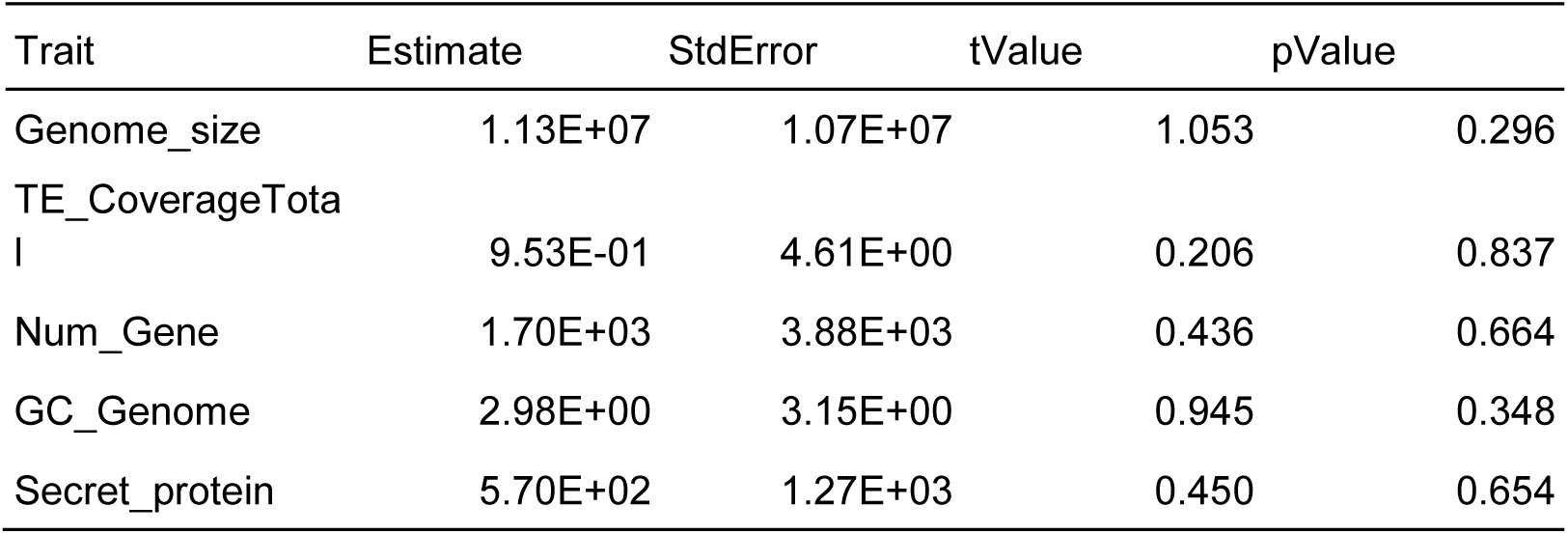
Phylogenetically Independent Contrasts (PIC) analysis results for genomic features in relation to ambrosia lifestyle. No significant associations were found after controlling for phylogeny

### Metabolic Versatility and Genomic Stability in Ambrosia Fungi

Our analysis of functional gene families in ambrosia fungi revealed a more straightforward evolutionary picture than previously thought. Unlike the extreme genome reduction often seen in many prokaryotic symbionts ^75,76^ or some eukaryotic symbionts ^77,78^, we observed nearly equivalent mean copy numbers of genes in ambrosia fungi (3.51) compared to non-ambrosia fungi (3.52). This maintenance of gene copy numbers aligns with observations in other eukaryotic symbionts, including some mycorrhizal fungi ^11,77^ and lichen mycobiont ^79^.

Specific gene families crucial for fungal metabolism differed between fungal clades (Figure 3A). CAZymes showed a slight increase in ambrosia fungi, with a gain/loss ratio of 1.12, while Plant Cell Wall Degrading Enzymes (PCWDEs) showed near-balance to slight reductions (gain/loss ratio of 0.89). This suggests adaptation to the utilization of simpler carbohydrates compared to complex wood polymer degradation in some free-living relatives. These findings indicate a refinement of these enzyme families, not a substantial decrease, contrasting with the loss of many biosynthetic pathways often seen in other symbionts ^69,78,80^.The near-balance to slight reduction in GPCRs (gain/loss ratio of 0.818) observed in ambrosia fungi likely reflects their dual nature as symbionts that retain many genetic and metabolic characteristics of their free-living relatives. Despite their symbiotic lifestyle within beetle galleries, these fungi still require mechanisms to sense and respond to environmental cues, much like their non-symbiotic counterparts. The absence of Pth11-like GPCRs, often associated with host-pathogen interactions, further suggests that ambrosia fungi have maintained environmental sensing capabilities, but our data do not allow to distinguish whether these GPCRs are related to interactions with the colonized host tree tissues, or with the mycangium of the vector beetle. Lipases and peptidases showed more widespread reductions, with gain/loss ratios of 0.495 and 0.418, respectively. However, these losses primarily involve genes with low copy numbers (Table S4). This pattern is reminiscent of gene family contractions observed in some eukaryotic symbionts, such as the yeast-like symbiont (YLS) of aphids, which maintains most metabolic pathways but shows some gene family reductions ^81^. The observed gene reduction may be an adaptive trait, a consequence of the ambrosia lifestyle evolution, or a result of genetic drift in small, clonal populations with limited sexual reproduction. Distinguishing between these possibilities remains challenging.

**Figure 3.**
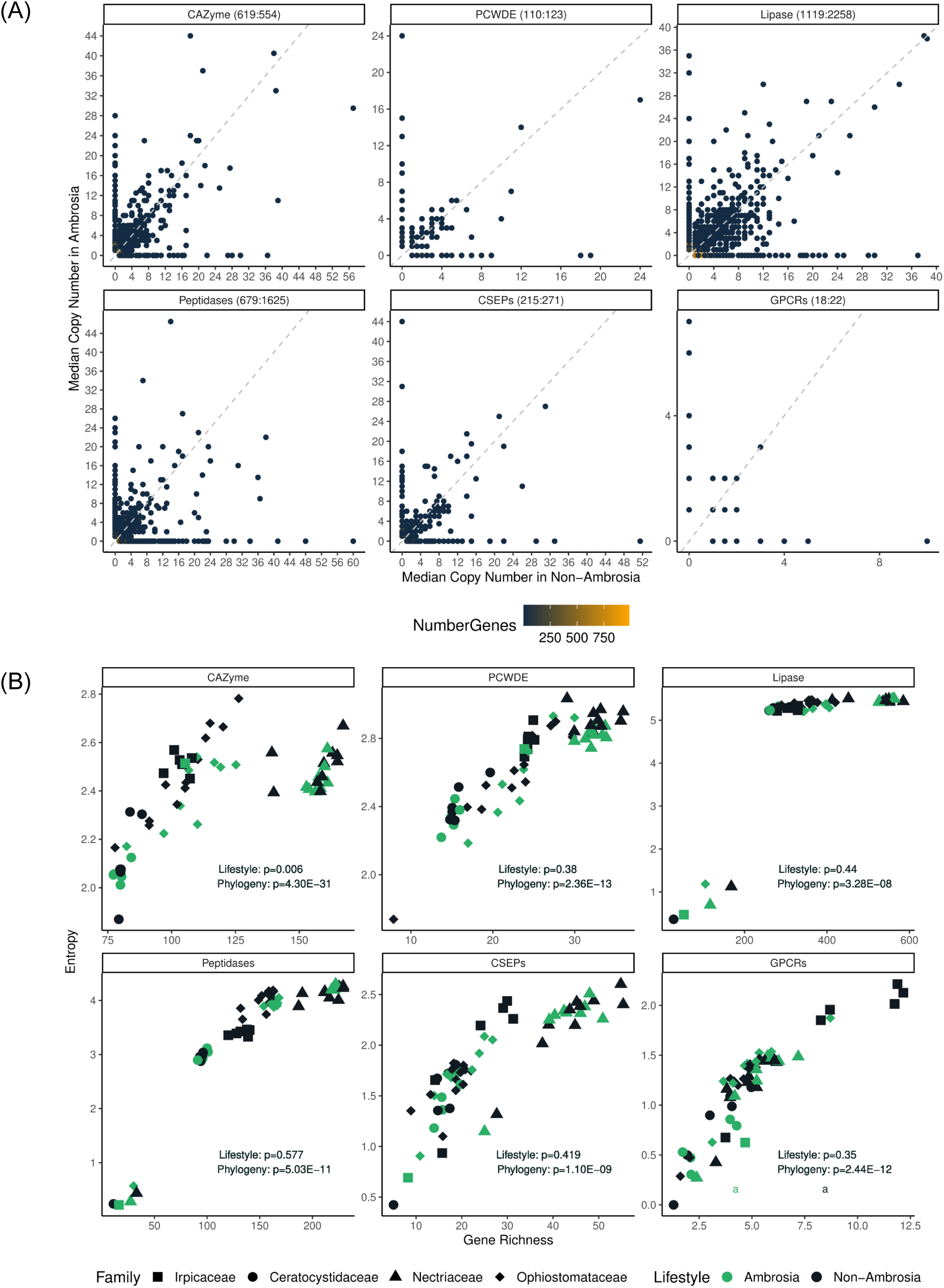
Gene repertoire dynamics in ambrosia fungi. (A) Gene copy number comparison between ambrosia and non-ambrosia fungi. Each point represents a gene family, with its position indicating mean copy numbers in ambrosia (y-axis) and non-ambrosia (x-axis) fungi. Color intensity reflects the number of genes at that ratio. Dashed line represents equal copy numbers. Inset values show total gains and losses per functional group in ambrosia fungi (gain:loss). (B) Functional diversity analysis. Each point represents a genome, with x-axis showing gene count and y-axis showing Shannon entropy diversity. Green dots indicate ambrosia fungi, black dots indicate non-ambrosia fungi. Point shapes denote fungal families: squares for Irpicaceae, circles for Ceratocystidaceae, triangles for Nectriaceae, and diamonds for Ophiostomataceae. P-values denote significance of diversity differences between ambrosia lifestyle (Lifestyle) or fungal family (Phylogeny) for each gene category.

A test of correlation between functional gene richness and diversity, expressed as Shannon entropy (Figure 3B) revealed a much greater influence of phylogenetic relatedness than the effect of ambrosial lifestyle. CAZymes was the only functional gene tested that showed significant difference (p = 0.006) in relation to lifestyle, indicating a potential impact of the ambrosia habit on carbohydrate metabolism. For other gene categories, including PCWDEs, lipases, peptidases, CSEPs, GPCRs, the lifestyle did not significantly influence functional diversity (p > 0.05). The gene richness data also revealed notable patterns, showing a consistently strong phylogenetic effect across all categories, with p-values ranging from 3.28E-08 to 4.30E-31. This variability in functional diversity aligns with the observation that eukaryotic symbionts often show more variable patterns of gene richness modification compared to prokaryotic symbionts ^12,81–83^.

The retention of genome size and metabolic diversity in ambrosia fungi likely relates to their largely free-living lifestyle. Unlike internal symbionts, ambrosia fungi face constant cycles of colonization and competition in the ephemeral moribund wood. This requires maintaining diverse metabolic capabilities ^3,15^ to quickly establish dominance, outcompete other fungi ^84^, and utilize varied resources. Moreover, the need to re-enter the mycangia of dispersing adult beetles adds another layer of selective pressure ^43,47^. Ambrosia fungi must not only dominate the gallery environment but also maintain their ability to grow in a form suitable for mycangial transport (Francke-Grosmann 1967). The genomic “refinement” of ambrosia fungi can be viewed as an adaptive strategy balancing metabolic versatility with symbiotic specialization. This evolutionary trajectory allows them to remain competitive while meeting symbiotic demands, demonstrating that obligate symbiosis doesn’t always lead to extreme genome reduction, especially in variable environments.

### Differential Evolution of CAZyme Families in Ambrosia Fungi

Our PIC analysis reveals family-specific enzyme evolution. Instead of observing a uniform trend across ambrosia lineages, we found distinct patterns of CAZyme family expansions and contractions within different fungal families, underscoring the necessity of considering phylogenetic relationships when interpreting these results (Table S5). In our overall analysis, we identified enzyme families such as AA4, CE4, and GH3 as having significant associations with the ambrosia lifestyle.

The AA4 family, which includes vanillyl-alcohol oxidases involved in lignin modification, showed a significant positive association with the ambrosia lifestyle in the PIC analysis (Estimate = 1.61, p = 0.011). Interestingly, this expansion is primarily driven by the ancestral non-ambrosial lineages within the Nectriaceae family, particularly in *Neocosmospora* species [*N. metavorans* (Al-Hatmi, S.A. Ahmed & de Hoog) Sand.-Den. & Crous, *N. liriodendra* Sand.-Den. & Crous, and *N. solani*). This interesting history suggests an expansion of the metabolic versatility followed by a contraction in a lineage adapted to insect-vectoring, but this scenario remains a hypothesis, as neither the timing of these events nor any comparable system are available.

The CE4 family, which includes acetyl xylan esterases involved in hemicellulose modification, showed a significant expansion in the PIC analysis (Estimate = 0.81, p = 0.005) in the Ophiostomataceae ambrosia lineages. This family-specific expansion suggests that enhanced hemicellulose modification capabilities may be especially important for Ophiostomataceae ambrosia fungi, possibly reflecting differences in wood substrate preferences or gallery construction strategies among different ambrosia symbioses. Similar to the findings of De Fine Licht and Biedermann ^85^, which were the first to report functional hemicellulolytic enzymes in ambrosia fungi.

The GH3 family, which includes β-glucosidases crucial for cellulose degradation, showed a significant expansion in the PIC analysis (Estimate = 0.65, p = 0.046). Notably, this expansion is more prominent in both Ophiostomataceae and Ceratocystidaceae ambrosia lineages. The expansion of GH3 in these two families suggests that enhanced cellulose degradation capabilities may be a shared adaptation in these groups of ambrosia fungi. This could indicate a convergent evolution towards more efficient cellulose processing,potentially allowing for better nutrient extraction from and manipulation of their ambient wood. The expansion of both CE4 and GH3 in Ophiostomataceae ambrosia fungi further suggests that this group may have evolved particularly enhanced capabilities for both hemicellulose and cellulose modification, supporting the conclusions of our enzyme activity analysis.

It is important to note that the signals from Irpicaceae ambrosia lineages are relatively weak in the overall PIC analysis, as only one genome from this group, generated in the present study, was included. This underscores the diversity of evolutionary strategies among ambrosia fungi and suggests that different fungal families may have adapted to the ambrosia lifestyle through distinct enzymatic modifications ^86^.

These results highlight the diverse nature of the ambrosia symbiosis across different fungal lineages.,The evolution of the CAZyme repertoire in ambrosia fungi varies significantly among fungal families. Rather than a uniform loss or gain of wood-degrading functions, it appears that ambrosia fungi have retained or evolved specialized enzymatic toolkits optimized for their unique ecological niches and phylogenetic backgrounds. The lineage-specific patterns of CAZyme evolution suggest that the transition to the ambrosia lifestyle may have occurred through different enzymatic adaptations in each fungal family. The expansion patterns of AA4, CE4, and GH3 families provide evidence for specialized adaptations in lignin modification, hemicellulose deacetylation, and cellulose degradation, respectively.

**Figure 4.**
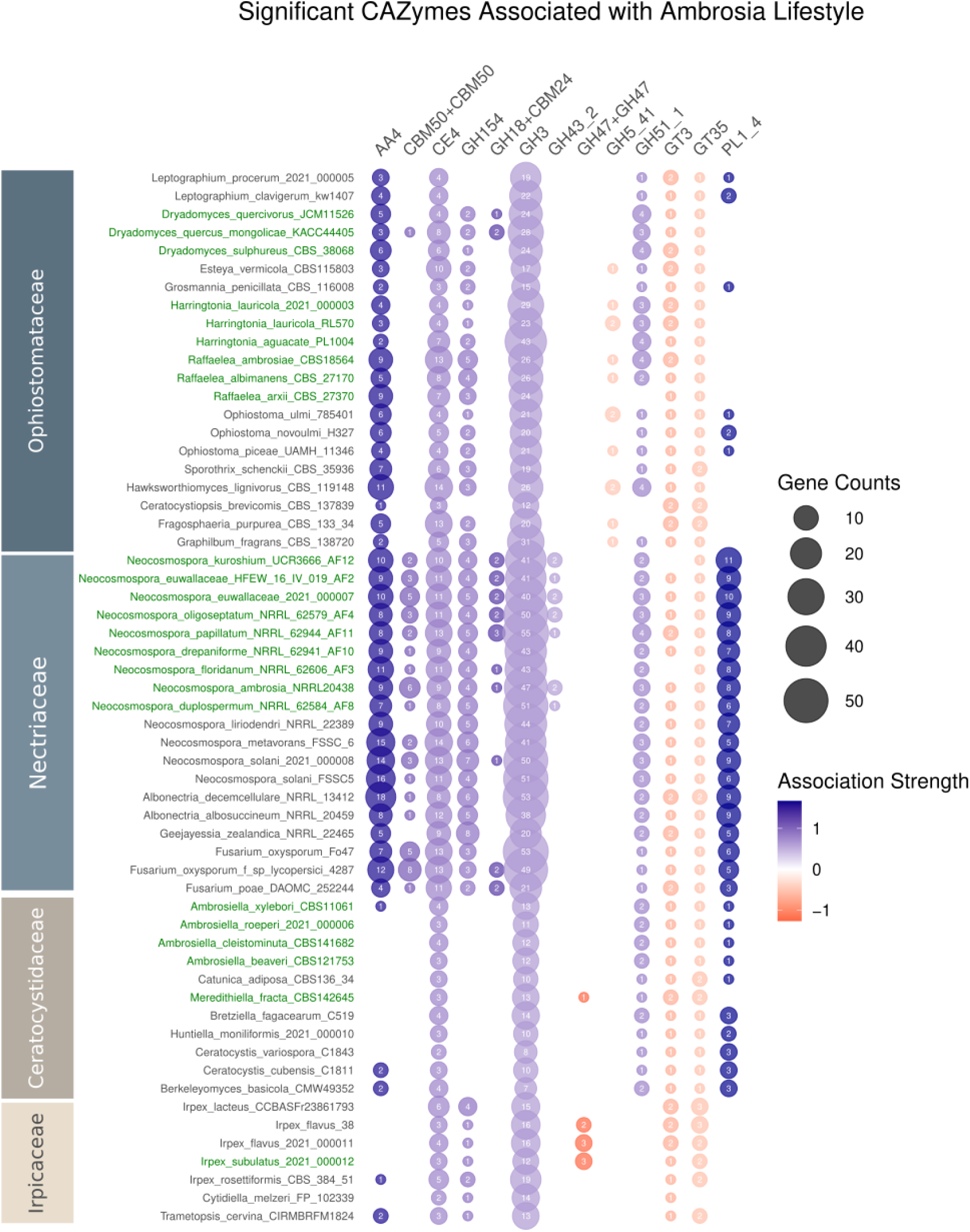
Bubble plot illustrating patterns of CAZyme enrichment and depletion across ambrosia fungi (green) and their non-ambrosia relatives (black). CAZyme families shown are significantly associated with ambrosia lifestyle (p < 0.05). Bubble size represents gene count per species. X-axis: CAZyme families, color-coded by association direction (blue: positive, red: negative), with color intensity indicating association strength. Y-axis: fungal species in the PIC test, ambrosia lineages are highlighted in green.

### RIP Activity and Genomic Adaptation in Ambrosia Fungi

Repeat-Induced Point (RIP) mutations do not differ overall between ambrosia and non-ambrosia fungi. Statistical analysis showed no difference in RIP-affected windows (Mann-Whitney U-test, p = 0.981), RIP-affected genomic proportion (p = 0.881), or count of Large RIP Affected Regions (LRARs) (p = 0.859). This lack of statistical difference suggests that ambrosia lifestyle alone is not a primary determinant of RIP activity.

Most fungi in our dataset possessed the key methyltransferases DIM-2 and RID, as well as several cofactors involved in DNA methylation and heterochromatin formation, regardless of their ecology. Only *Dryadomyces quercus-mongolicae* and *D. sulphureus* lacked a detectable RID gene, while the free-living relative *Hawksworthiomyces lignivorus* lacked both RID and HP-1. The retention of these genes across most species, regardless of their ambrosia status or observed RIP activity, suggests that factors beyond symbiotic lifestyle influence the maintenance of these genomic elements. However, the presence of these genes alone does not guarantee RIP activity ^87^. Their maintenance in ambrosia fungi could indicate potential for cryptic sexual cycles ^32,33^, additional gene functions beyond repeat silencing, or recent loss of RIP activity with insufficient time for gene loss.

**Figure 5.**
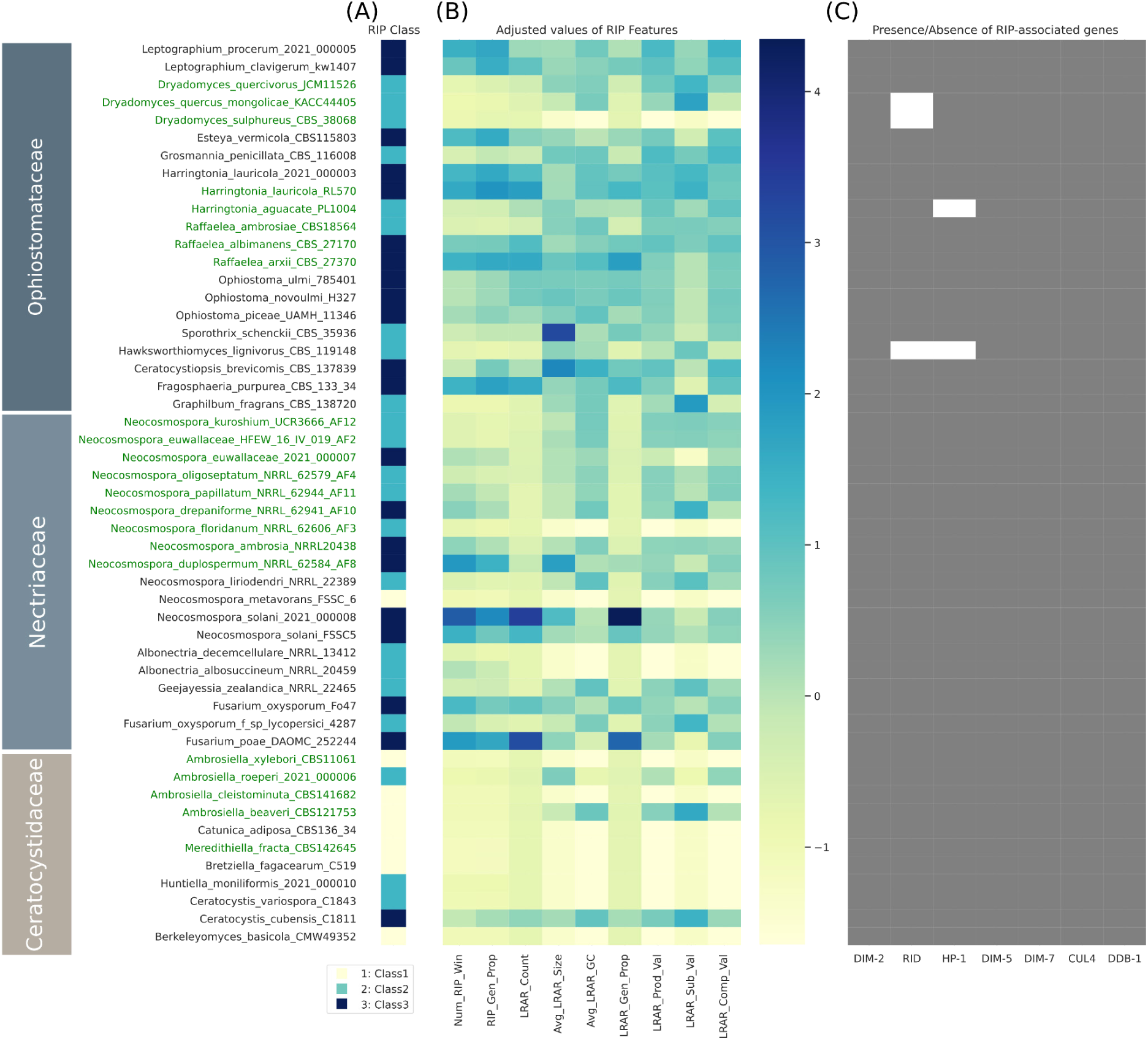
Genome-wide RIP analyses of ambrosia (green) and non-ambrosia (black) fungi. (A) RIP classes assigned based on the proportion of the genome affected by RIP, ranging from Class 1 (0–0.2%) to Class 3 (1–5%) ^88^. (B) Heatmap of standardized values for key RIP features: Number of RIP Affected Windows (Num_RIP_Win), RIP Affected Genomic Proportion (%) (RIP_Gen_Prop), Count of LRARs (LRAR_Count), Average Size of LRARs (bp) (Avg_LRAR_Size), Average GC Content of LRARs (%) (Avg_LRAR_GC), Genomic Proportion of LRARs (bp) (LRAR_Gen_Prop), Product Value for LRARs (LRAR_Prod_Val), Substrate Value for LRARs (LRAR_Sub_Val), and Composite Value for LRARs (LRAR_Comp_Val). (C) Presence (gray) or absence (white) of RIP-associated genes identified by blast against *Neurospora crassa* Shear & B.O. Dodge homologs. Irpicaceae species were excluded as RIP is primarily observed in Ascomycota.

### Conclusions

These findings underscore the diverse molecular ecology of the role of ambrosia fungi in wood, as well as their interactions with their insect vectors. Adaptations toward ambrosia symbiosis may be more diverse and sophisticated than previously recognized, with different lineages adopting distinct strategies. Given the lineage-specific evolution of fungal metabolism, future research should focus on the biochemical characterization of these enzyme families within specific fungal lineages, while harnessing the power of comparative analysis that is so unique to the ambrosia symbiosis. Future improved and more detailed assemblies of fungal genomes will also improve our capacity to study the fine-scale adaptation to the symbiotic lifestyle which were not available in our analysis, such as the mating genes to determine the evolution of sexuality in the ambrosia fungi, and especially the genetic basis of the interactions between the fungi and the beetles. Experimental data, like transcriptomes sampled during critical periods of the relationships’ lifecycle, may be necessary to infer the evolution of mutual signaling of nutrition provisioning, which may be key to our understanding of the crux of the relationship — the mechanisms maintaining mutual specificity to the exclusion of other organisms — and their roles in different ambrosia beetle-fungus symbioses to fully understand their functional significance in these fascinating ecological relationships.

## Material and Methods

### Fungal Strain Selection and Preparation

The fungal strains used in this study (Table S1) represent a spectrum of ecological associations within the ambrosia beetle system. Ambrosia fungi were defined as dominant and consistent mycangial isolates. In the analyses, we paired these with non-ambrosia counterparts selected based on phylogenetic relatedness and ecological relevance. The non-ambrosia fungi were isolated either from ambrosia beetle body surfaces or from associated wood substrates. This approach allowed for comparative genomic analysis across different degrees of symbiotic association within similar ecological niches.

### DNA Extraction and Quality Assessment

DNA was extracted from 10-day-old fungal cultures. Fungal tissues were scraped from PDA medium and placed into 1.5 ml microcentrifuge tubes containing 200 mg zirconia beads. The tubes were shaken using a Vortex-Genie 2 (Scientific Industries, Inc., NY, USA) at 3200 rpm for 4 minutes to disrupt the mycelium. DNA purification was carried out using a modified Bio-On-Magnetic-Beads (BOMB) protocol (Oberacker et al. 2019) with the ZiXpress 32 automated nucleic acid purification instrument (New Taipei City, Taiwan). DNA quality was assessed using an Invitrogen Qubit Fluorometer (ThermoFisher Scientific).

### Genome Sequencing, Assembly, and Quality Assessment

Genomic DNA was sequenced using Oxford Nanopore Technology. Library preparation was performed using the Native Barcoding Kit 24 V14, following the manufacturer’s protocol. We loaded 0.4 μg of DNA library onto an R10.4.1 flow cell and sequenced on a MinION device using MinKNOW software v23.07.5. Raw sequence outputs were basecalled using dorado v0.5.1 (github.com/nanoporetech/dorado) with the SUP model. Sequences were then processed using the nanoACT pipeline (github.com/Raingel/nanoACT). This processing included quality-filtering (q-score = 9, minimum length = 500, and maximum length = 1,000,000), demultiplexing using singlebar(), and trimming using trim_reads(). Trimmed reads were de novo assembled using Flye v2.9.3 with default settings and the "--nano-hq" option. Consensus sequences of the assemblies were polished using Medaka v1.8.1 https://github.com/nanoporetech/medaka) with Oxford nanopore reads. Transposable element (TE) coverage in the genomes was identified using RepeatModeler v2.0.5 and RepeatMasker v4.1.6. Genome completeness was assessed using BUSCO v5.6.1 with fungi-odb10 as a reference and default parameters. The resulting genome assemblies were deposited in NCBI under Bioproject accession PRJNA1084195 (Table S1).

### Comparative genomic feature analyses

We performed genome annotation using BRAKER3 v3.0.5, which integrates RNA-Seq data from the Sequence Read Archive (SRA) with gene predictions from AUGUSTUS and GeneMark-ET. Gene functions were subsequently assigned using eggNOG-mapper v2.1.12 with the eggNOG orthology v5.0.2 database.

To predict the fungal secretome, we annotated our genomes using multiple databases. We identified signal peptides using SignalP 6.0 ^89^, transmembrane domains using TMHMM via Galaxy (Krogh et al. 2001; Afgan et al. 2018), and subcellular localization using TargetP (Armenteros and Salvatore 2019). Proteins meeting any of these criteria were included in the predicted secretome. We further characterized the secretome for lipases and peptidases by aligning protein sequences to the Lipase Engineering Database (LED) ^90^ and MEROPS peptidase database ^91^ using DIAMOND v2.1.8 ^92^. The top hit for each protein was selected based on bitscore. Carbohydrate-Active Enzymes (CAZymes) were annotated using dbCAN3 ^93^ with HMMER, Diamond, and dbCAN_sub for CAZyme families annotation. Specifically, we only included annotated CAZymes genes that were concurrently annotated by both HMMER and dbCAN_sub. We categorized CAZymes into Plant Cell Wall-Degrading Enzymes (PCWDEs) following the classification scheme of Miyauchi et al. (2020). G-protein coupled receptors (GPCRs) were identified through a BLASTP search against the best hits in the GPCRDB database ^94^, and their functional roles were cross-referenced with UniProt ^95^. To identify CSEPs, we used Effectorp 3.0. We retained both apoplastic and cytoplasmic effectors in our analysis, excluding only those identified as "non-effector" by the tool. All analyses were performed using default parameters unless otherwise specified.

We performed Multivariate Analysis of Variance (MANOVA) to examine the effects of ambrosia lifestyle and fungal family on diversity indices across various genomic features. The analysis was conducted using the manova() in R. For each functional gene category, we carried out separate MANOVAs to assess the impact of ambrosia lifestyle and family on two diversity indices: gene richness and Shannon entropy. The Pillai’s trace statistic was used to evaluate the significance of the effects.

Key genomic features were compared between ambrosia and non-ambrosia fungi: genome size, transposable element (TE) coverage, number of genes, scaffold N50, number of secreted proteins, and genomic GC content. To control for phylogenetic non-independence, we performed Phylogenetically Independent Contrasts (PIC) analysis using the ape ^96^ and phangorn ^97^ R packages. This method calculates contrasts in trait values between phylogenetically related species, accounting for shared evolutionary history. Linear models were then fitted to these contrasts, with genomic features as dependent variables and ambrosia lifestyle as the independent variable. These models quantify the association between each genomic feature and the ambrosia lifestyle while controlling for phylogeny, providing statistical estimates for each trait-lifestyle relationship.

### Repeat-Induced Point Mutations Estimation

Repeat masked whole-genome sequences of ambrosia and non-ambrosia fungi were used to estimate Repeat-Induced Point (RIP) mutations in their genomes. RIP mutations were detected using The RIPper software ^98^ with stringent RIP parameters (RIP product > 1.1, RIP substrate ≤ 0.75, and RIP composite index > 7). The analysis employed a sliding-window approach with 1,000 bp windows and 500 bp step sizes.

RIP-associated genes (DIM-2, RID, HP-1, DIM-5, DIM-7, CUL4, and DDB-1) were identified using BLASTp searches against these genes identified in *Neurospora crassa* (XP_959891.1, XP_957632.1, XP_011392925.1, XP_957479.2, XP_961308.2, XP_962347.1, and XP_957743.3, respectively), with an E-value cutoff of 1e-5. The Mann-Whitney U test was used to compare RIP-related metrics between ambrosia and non-ambrosia fungi. Four key features were analyzed: number of RIP-affected windows, RIP-affected genomic proportion, and count of Large RIP Affected Regions (LRARs). Analyses were conducted both across all fungi and within individual fungal families.

Heatmaps were generated using seaborn and matplotlib libraries in Python. The heatmaps visualized RIP class, standardized RIP features, and gene presence/absence across fungal species. RIP classes were assigned based on the proportion of the genome affected by RIP, ranging from Class 1 (0-0.2%) to Class 6 (≥20%) according to ^88^. RIP features were standardized using sklearn’s StandardScaler.

### Phylogenetic Reconstruction

We used representative genomes of fungi across Basidiomycota and Ascomycota, including the most closely-related species to ambrosia fungi, along with two Mucoromycota representatives as outgroups, to reconstruct phylogenetic relationships. BUSCO v5.5.0_cv1 was run with the fungi_odb10 lineage database to generate single-copy orthologs from each genome. The shared single-copy orthologs were used as input for building the species tree. Sequences were aligned individually using MAFFT v7.490 with default parameters. The alignments were then trimmed using ClipKIT v2.3.0 with the "smartgap" option. We retained BUSCO gene alignments with ≥90% taxon occupancy for further analysis. Output alignments from ClipKIT were used as input to ModelTest-NG v0.1.7 with default settings to select the best evolutionary model for each gene. For the species tree reconstruction, we used a concatenation-based method. The set of retained 742 BUSCO gene alignments from MAFFT and ClipKIT were concatenated. The alignment was partitioned by gene, using the best evolutionary model as predicted with ModelTest-NG. We reconstructed the tree using IQ-TREE2 v2.0.7 with parameters "-bb 1000 -alrt 1000." The resulting tree was rooted to the Mucoromycota outgroups in FigTree. For the gene tree reconstruction, we analyzed each single-copy ortholog alignment separately. The same set of BUSCO gene alignments were processed using IQ-TREE2 with the evolutionary models predicted by ModelTest-NG, ensuring each gene tree was accurately reconstructed. All resulting gene trees were concatenated into a single file. ASTRAL v5.7.1 ^99^ was then employed to infer individual phylogenies from these BUSCO genes.

To test for significant discordance between gene trees and the species tree, we performed chi-square tests comparing the distribution of concordant and discordant gene tree topologies to the expectation under incomplete lineage sorting (ILS). For each branch in the species tree, we calculated the gene concordance factor (gCF) and site concordance factor (sCF) using IQ-TREE. We then conducted chi-square tests comparing the observed number of concordant vs discordant genes (for gCF) and sites (for sCF) to the expected 1/3 ratio under ILS. A custom R function was used to perform these tests for each branch, with p-values < 0.05 considered significant evidence of discordance beyond ILS expectations. Branches showing significant discordance in either gCF or sCF were identified as potentially resulting from processes other than ILS, such as introgression or incomplete taxon sampling.

### Divergence Dating

To estimate the timing of ambrosial domestication events, we employed the RelTime method in MEGA X ^100^, which calculates relative divergence times by leveraging molecular evolutionary rates across various branches of the phylogenetic tree ^101,102^. We input the concatenated alignments of single-copy BUSCO orthologs and the maximum likelihood (ML) topologies generated from our phylogenetic reconstruction. Our analysis incorporated three calibration nodes based on fossil records and the TimeTree database ^103^: the divergence between Ascomycota and Basidiomycota (582 to 749 Million years ago [Ma]), the split between Sclerotiniaceae and Sordariomycetes (239.7 -320 Ma), and a fossil record constraining the minimum age of Diaporthales (> 136 Ma) ^104^. We modeled the molecular data using the Tamura-Nei model with gamma-distributed rates among sites (TN93+G), which was selected as the best fit based on Bayesian Information Criterion (BIC) scores.

### Ancestral State Reconstruction

We performed ancestral state reconstruction (ASR) to infer the evolutionary history of ambrosia status on the 61 taxa phylogeny inferred from our concatenated supermatrix dataset. The analysis was conducted using a maximum likelihood approach and stochastic character mapping. We used the ape and phytools ^105^ packages in R for these analyses.

First, we used the ace() in the ape package to perform maximum likelihood ancestral state reconstruction. This method estimates the marginal ancestral state likelihoods for all nodes within the phylogeny, assuming that the probability of character state changes depends only on the current state and not on previous states. We applied the default equal rates (ER) model, which assumes equal transition rates between all character states. To account for uncertainty in the ancestral state estimates, we also conducted stochastic character mapping using the make.simmap function from the phytools package. This approach uses Markov chain Monte Carlo (MCMC) to sample character histories from their posterior probability distribution. We generated 1000 simulated character histories on the phylogeny using the ER model. The results of these simulations were summarized to obtain posterior probabilities of ancestral states at each node.

We visualized the results of both the maximum likelihood reconstruction and the stochastic character mapping on the phylogenetic tree. For the maximum likelihood results, we plotted pie charts at each internal node representing the probability of each ancestral state. For the stochastic character mapping results, we plotted a summary of the 1000 simulated character histories, with pie charts at nodes showing the proportion of simulations supporting each state.

## Acknowledgements

YTH, KA, and GJP was supported by the National Science and Technology Council, Taiwan (110-2621-B-037-002-MY3). JHO was supported by the National Agriculture and Food Research Organization, Japan. JH was supported by the US National Science Foundation and the USDA Forest Service.

---

**Figure S1.**
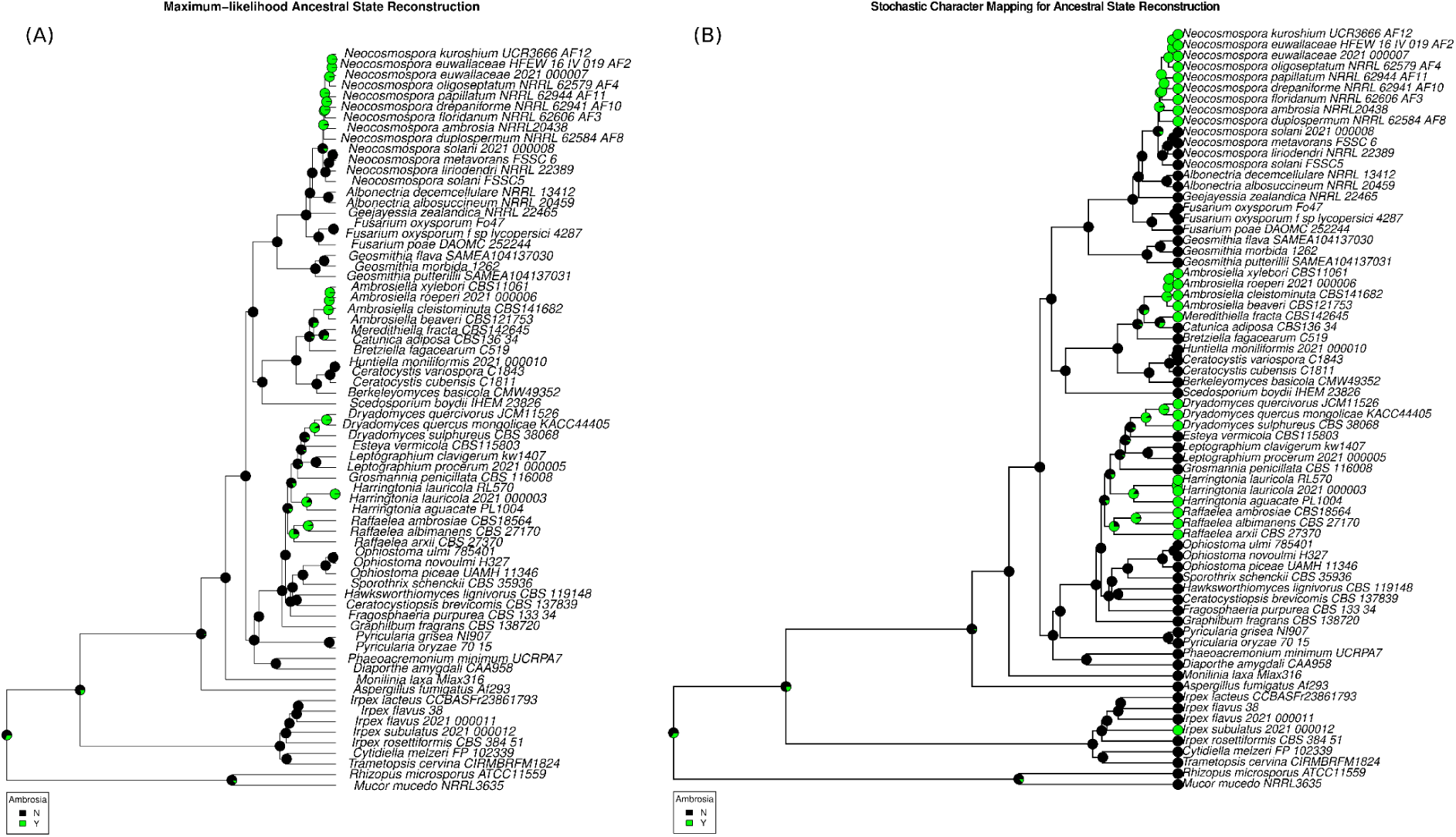
Ancestral State Reconstruction of Ambrosia Fungi. (A) Maximum-Likelihood Method. (B) Stochastic Character Mapping Method.

## Reference

1. Francke-Grosmann, H. Some New Aspects in Forest Entomology. Annu. Rev. Entomol. 8, 415–438 (1963).

2. Hulcr, J. & Stelinski, L. L. The Ambrosia Symbiosis: From Evolutionary Ecology to Practical Management. Annu. Rev. Entomol. 62, 285–303 (2017).

3. Huang, Y.-T., Skelton, J. & Hulcr, J. Lipids and small metabolites provisioned by ambrosia fungi to symbiotic beetles are phylogeny-dependent, not convergent. ISME J. 14, 1089–1099 (2020).

4. Harrington, T. C., McNew, D., Mayers, C., Fraedrich, S. W. & Reed, S. E. Ambrosiella roeperi sp. nov. is the mycangial symbiont of the granulate ambrosia beetle, Xylosandrus crassiusculus. Mycologia 106, 835–845 (2014).

5. Freeman, S. et al. Fusarium euwallaceae sp. nov.--a symbiotic fungus of Euwallacea sp., an invasive ambrosia beetle in Israel and California. Mycologia 105, 1595–1606 (2013).

6. Kirkendall, L. R., Biedermann, P. H. W. & Jordal, B. H. Evolution and Diversity of Bark and Ambrosia Beetles. in Bark Beetles (eds. Vega, F. E. & Hofstetter, R. W.) 85–156 (Academic Press, San Diego, 2015). doi:10.1016/B978-0-12-417156-5.00003-4.

7. Johnson, A. J. et al. Phylogenomics clarifies repeated evolutionary origins of inbreeding and fungus farming in bark beetles (Curculionidae, Scolytinae). Mol. Phylogenet. Evol. 127, 229–238 (2018).

8. Pistone, D., Gohli, J. & Jordal, B. H. Molecular phylogeny of bark and ambrosia beetles (Curculionidae: Scolytinae) based on 18 molecular markers. Syst. Entomol. 43, 387– 406 (2018).

9. Li, Y. et al. New Fungus-Insect Symbiosis: Culturing, Molecular, and Histological Methods Determine Saprophytic Polyporales Mutualists of Ambrosiodmus Ambrosia Beetles. PLoS One 10, e0137689 (2015).

10. Massoumi Alamouti, S., Tsui, C. K. M. & Breuil, C. Multigene phylogeny of filamentous ambrosia fungi associated with ambrosia and bark beetles. Mycol. Res. 113, 822–835 (2009).

11. Miyauchi, S. et al. Large-scale genome sequencing of mycorrhizal fungi provides insights into the early evolution of symbiotic traits. Nat. Commun. 11, 5125 (2020).

12. Nygaard, S. et al. Reciprocal genomic evolution in the ant-fungus agricultural symbiosis. Nat. Commun. 7, 12233 (2016).

13. Fu, N., Wang, M., Wang, L., Luo, Y. & Ren, L. Genome Sequencing and Analysis of the Fungal Symbiont of Sirex noctilio, Amylostereum areolatum: Revealing the Biology of Fungus-Insect Mutualism. mSphere 5, (2020).

14. Poulsen, M. et al. Complementary symbiont contributions to plant decomposition in a fungus-farming termite. Proc. Natl. Acad. Sci. U. S. A. 111, 14500–14505 (2014).

15. Huang, Y.-T., Skelton, J. & Hulcr, J. Multiple evolutionary origins lead to diversity in the metabolic profiles of ambrosia fungi. Fungal Ecol. 38, 80–88 (2019).

16. Lehenberger, M., Biedermann, P. H. W. & Benz, J. P. Molecular identification and enzymatic profiling of Trypodendron (Curculionidae: Xyloterini) ambrosia beetle-associated fungi of the genus Phialophoropsis (Microascales: Ceratocystidaceae). Fungal Ecol. 38, 89–97 (2019).

17. Veselská, T. et al. Adaptive traits of bark and ambrosia beetle-associated fungi. Fungal Ecol. 41, 165–176 (2019).

18. Resl, P. et al. Large differences in carbohydrate degradation and transport potential among lichen fungal symbionts. Nat. Commun. 13, 2634 (2022).

19. Hernández-Martínez, F., Briones-Roblero, C. I., Nelson, D. R., Rivera-Orduña, F. N. & Zúñiga, G. Cytochrome P450 complement (CYPome) of Candida oregonensis, a gut-associated yeast of bark beetle, Dendroctonus rhizophagus. Fungal Biol. 120, 1077– 1089 (2016).

20. Skelton, J. et al. Fungal symbionts of bark and ambrosia beetles can suppress decomposition of pine sapwood by competing with wood-decay fungi. Fungal Ecol. 45, 100926 (2020).

21. Brown, N. A., Schrevens, S., van Dijck, P. & Goldman, G. H. Fungal G-protein-coupled receptors: mediators of pathogenesis and targets for disease control. Nat. Microbiol. 3, 402–414 (2018).

22. Xue, C., Hsueh, Y.-P. & Heitman, J. Magnificent seven: roles of G protein-coupled receptors in extracellular sensing in fungi. FEMS Microbiol. Rev. 32, 1010–1032 (2008).

23. Kou, Y., Tan, Y. H., Ramanujam, R. & Naqvi, N. I. Structure-function analyses of the Pth11 receptor reveal an important role for CFEM motif and redox regulation in rice blast. New Phytol. 214, 330–342 (2017).

24. Shang, J., Shang, Y., Tang, G. & Wang, C. Identification of a key G-protein coupled receptor in mediating appressorium formation and fungal virulence against insects. Sci. China Life Sci. 64, 466–477 (2021).

25. Plett, J. M. et al. A secreted effector protein of Laccaria bicolor is required for symbiosis development. Curr. Biol. 21, 1197–1203 (2011).

26. Martin, F., Kohler, A., Murat, C., Veneault-Fourrey, C. & Hibbett, D. S. Unearthing the roots of ectomycorrhizal symbioses. Nat. Rev. Microbiol. 14, 760–773 (2016).

27. Garcia, K. & Ané, J.-M. Comparative analysis of secretomes from ectomycorrhizal fungi with an emphasis on small-secreted proteins. Front. Microbiol. 7, 1734 (2016).

28. Farrell, B. D. et al. The evolution of agriculture in beetles (Curculionidae: Scolytinae and Platypodinae). Evolution 55, 2011–2027 (2001).

29. Normark, B. B., Judson, O. P. & Moran, N. A. Genomic signatures of ancient asexual lineages. Biol. J. Linn. Soc. Lond. 79, 69–84 (2003).

30. Harrington, T. C., Aghayeva, D. N. & Fraedrich, S. W. New combinations in Raffaelea, Ambrosiella, and Hyalorhinocladiella, and four new species from the redbay ambrosia beetle, Xyleborus glabratus. Mycotaxon 111, 337–361 (2010).

31. Harrington, T. C. Ecology and evolution of mycophagous bark beetles and their fungal partners. in Vega F.E. *and* Blackwell, M. *(eds)* Insect-fungal associations: ecology and evolution (Oxford University Press: 257–291, 2005).

32. Mayers, C. G., Harrington, T. C. & Ranger, C. M. First report of a sexual state in an ambrosia fungus: Ambrosiella cleistominuta sp. nov. associated with the ambrosia beetle Anisandrus maiche. Botany 95, 503–512 (2017).

33. Mayers, C. G. et al. Patterns of coevolution between ambrosia beetle mycangia and the Ceratocystidaceae, with five new fungal genera and seven new species. Persoonia 44, 41–66 (2020).

34. Nel, W. J., Wingfield, M. J., de Beer, Z. W. & Duong, T. A. Ophiostomatalean fungi associated with wood boring beetles in South Africa including two new species. Antonie Van Leeuwenhoek 114, 667–686 (2021).

35. Vanderpool, D., Bracewell, R. R. & McCutcheon, J. P. Know your farmer: Ancient origins and multiple independent domestications of ambrosia beetle fungal cultivars. Mol. Ecol. 27, 2077–2094 (2018).

36. de Beer, Z. W., Procter, M., Wingfield, M. J., Marincowitz, S. & Duong, T. A. Generic boundaries in the Ophiostomatales reconsidered and revised. Stud. Mycol. 101, 57–120 (2022).

37. Degnan, J. H. & Rosenberg, N. A. Gene tree discordance, phylogenetic inference and the multispecies coalescent. Trends Ecol. Evol. 24, 332–340 (2009).

38. Kolaczkowski, B. & Thornton, J. W. Performance of maximum parsimony and likelihood phylogenetics when evolution is heterogeneous. Nature 431, 980–984 (2004).

39. Jordal, B. H. Molecular phylogeny and biogeography of the weevil subfamily Platypodinae reveals evolutionarily conserved range patterns. Mol. Phylogenet. Evol. 92, 294–307 (2015).

40. Shin, S. et al. Phylogenomic data yield new and robust insights into the phylogeny and evolution of weevils. Mol. Biol. Evol. 35, 823–836 (2018).

41. Nakashima. Several types of the mycetangia found in platypodid ambrosia beetles (Coleoptera: Platypodidae). *matsumurana*. New series: journal of the … (1975).

42. Li, Y. et al. Specific and promiscuous ophiostomatalean fungi associated with Platypodinae ambrosia beetles in the southeastern United States. Fungal Ecol. 35, 42– 50 (2018).

43. Mayers, C. G., Harrington, T. C. & Biedermann, P. H. W. Mycangia define the diverse ambrosia beetle–fungus symbioses. in The Convergent Evolution of Agriculture in Humans and Insects (The MIT Press, 2022). doi:10.7551/mitpress/13600.003.0013.

44. Hulcr, J. & Skelton, J. Ambrosia beetles. in Forest Entomology and Pathology 339–360 (Springer International Publishing, Cham, 2023). doi:10.1007/978-3-031-11553-0_11.

45. Six, D. L. Ecological and Evolutionary Determinants of Bark Beetle -Fungus Symbioses. Insects 3, 339–366 (2012).

46. Gohli, J. et al. Biological factors contributing to bark and ambrosia beetle species diversification. Evolution 71, 1258–1272 (2017).

47. Skelton, J. et al. A selective fungal transport organ (mycangium) maintains coarse phylogenetic congruence between fungus-farming ambrosia beetles and their symbionts. Proc. Biol. Sci. 286, 20182127 (2019).

48. Francke-Grosmann, H. Ectosymbiosis in wood-inhabiting insects. Symbiosis 2, 141–205 (1967).

49. Greco, E. B. & Wright, M. G. Ecology, biology, and management of Xylosandrus compactus (Coleoptera: Curculionidae: Scolytinae) with emphasis on coffee in Hawaii. J. Integr. Pest Manag. 6, 7–7 (2015).

50. Galko, J. et al. Distribution, Habitat Preference, and Management of the Invasive Ambrosia Beetle Xylosandrus germanus (Coleoptera: Curculionidae, Scolytinae) in European Forests with an Emphasis on the West Carpathians. For. Trees Livelihoods 10, 10 (2018).

51. Storer, C., Payton, A., McDaniel, S., Jordal, B. & Hulcr, J. Cryptic genetic variation in an inbreeding and cosmopolitan pest, Xylosandrus crassiusculus, revealed using ddRADseq. Ecol. Evol. 7, 10974–10986 (2017).

52. Ranger, C. M., Reding, M. E., Addesso, K., Ginzel, M. & Rassati, D. Semiochemical-mediated host selection by*Xylosandrus*spp. ambrosia beetles (Coleoptera: Curculionidae) attacking horticultural tree crops: a review of basic and applied science. Can. Entomol. 153, 103–120 (2021).

53. Mayers, C. G., Bateman, C. C. & Harrington, T. C. New Meredithiella species from mycangia of Corthylus ambrosia beetles suggest genus-level coadaptation but not species-level coevolution. Mycologia 110, 63–78 (2018).

54. O’Donnell, K. et al. Discordant phylogenies suggest repeated host shifts in the Fusarium-Euwallacea ambrosia beetle mutualism. Fungal Genet. Biol. 82, 277–290 (2015).

55. Wang, Z. et al. The infestation and habitat of the ambrosia beetle Euwallacea interjectus (Coleoptera: Curculionidae: Scolytinae) in the riparian zone of Shanghai, China. Agric. For. Entomol. 23, 104–109 (2021).

56. Kasson, M. T. et al. An inordinate fondness for *Fusarium*: phylogenetic diversity of fusaria cultivated by ambrosia beetles in the genus *Euwallacea* on avocado and other plant hosts. Fungal Genet. Biol. 56, 147–157 (2013).

57. Carrillo, J. D. et al. Members of the Euwallacea fornicatus species complex exhibit promiscuous mutualism with ambrosia fungi in Taiwan. Fungal Genet. Biol. 133, 103269 (2019).

58. Hulcr, J., Black, A., Prior, K., Chen, C.-Y. & Li, H.-F. Studies of ambrosia beetles (Coleoptera: Curculionidae) in their native ranges help predict invasion impact. Fla. Entomol. 100, 257–261 (2017).

59. M Smith, S., F Gomez, D., A Beaver, R., Hulcr, J. & I Cognato, A. Reassessment of the Species in the Euwallacea Fornicatus (Coleoptera: Curculionidae: Scolytinae) Complex after the Rediscovery of the ‘Lost’ Type Specimen. Insects 10, 261 (2019).

60. Mendel, Z. et al. What determines host range and reproductive performance of an invasive ambrosia beetle Euwallacea fornicatus; Lessons from Israel and California. Front. For. Glob. Chang. 4, 654702 (2021).

61. Liao, Y.-C. et al. The Euwallacea fornicatus species complex (Coleoptera: Curculionidae); emerging economic pests of tea in Taiwan. Crop Prot. 168, 106226 (2023).

62. Jusino, M. A., et al. Relationships in Decay: Ambrosia Beetles Host Phylogenetically Diverse Basidiomycete Fungi. in (2020).

63. Jusino, M. A., Skelton, J., Chen, C.-C., Hulcr, J. & Smith, M. E. Sexual reproduction and saprotrophic dominance by the ambrosial fungus Flavodon subulatus (= Flavodon ambrosius). Fungal Ecol. 47, 100979 (2020).

64. Aanen, D. K. et al. The evolution of fungus-growing termites and their mutualistic fungal symbionts. Proc. Natl. Acad. Sci. U. S. A. 99, 14887–14892 (2002).

65. Inch, S., Ploetz, R., Held, B. & Blanchette, R. Histological and anatomical responses in avocado, Persea americana, induced by the vascular wilt pathogen, Raffaelea lauricola. Botany 90, 627–635 (2012).

66. Slippers, B., Wingfield, B. D., Coutinho, T. A. & Wingfield, M. J. DNA sequence and RFLP data reflect geographical spread and relationships of Amylostereum areolatum and its insect vectors. Mol. Ecol. 11, 1845–1854 (2002).

67. Hajek, A. E., Nielsen, C., Kepler, R. M., Long, S. J. & Castrillo, L. Fidelity among Sirex woodwasps and their fungal symbionts. Microb. Ecol. 65, 753–762 (2013).

68. Gomez, D. F. & Hulcr, J. The Punky Wood Ambrosia Beetle and Fungus in Florida that Cause Wood Rot: Ambrosiodmus minor and Flavodon subulatus: FOR365/FR434, 12/2020. Edis 2021, 4–4 (2021).

69. Kohler, A. et al. Convergent losses of decay mechanisms and rapid turnover of symbiosis genes in mycorrhizal mutualists. Nat. Genet. 47, 410–415 (2015).

70. Tian, C. F. et al. Comparative genomics of rhizobia nodulating soybean suggests extensive recruitment of lineage-specific genes in adaptations. Proc. Natl. Acad. Sci. U. S. A. 109, 8629–8634 (2012).

71. Smith, T. E., Li, Y., Perreau, J. & Moran, N. A. Elucidation of host and symbiont contributions to peptidoglycan metabolism based on comparative genomics of eight aphid subfamilies and their Buchnera. PLoS Genet. 18, e1010195 (2022).

72. Hulcr, J. et al. Mycangia of ambrosia beetles host communities of bacteria. Microb. Ecol. 64, 784–793 (2012).

73. Ibarra-Juarez, L. A. et al. Impact of rearing conditions on the ambrosia beetle’s microbiome. Life (Basel) 8, 63 (2018).

74. Kostovcik, M. et al. The ambrosia symbiosis is specific in some species and promiscuous in others: evidence from community pyrosequencing. ISME J. 9, 126–138 (2015).

75. van Ham, R. C. H. J. et al. Reductive genome evolution in *Buchnera aphidicola*. Proc. Natl. Acad. Sci. U.S.A. 100, 581–586 (2003).

76. Nakabachi, A. et al. The 160-kilobase genome of the bacterial endosymbiont Carsonella. Science 314, 267 (2006).

77. Martin, F. et al. Périgord black truffle genome uncovers evolutionary origins and mechanisms of symbiosis. Nature 464, 1033–1038 (2010).

78. Song, H. et al. A comparative genomic analysis of lichen-forming fungi reveals new insights into fungal lifestyles. Sci. Rep. 12, 10724 (2022).

79. Armaleo, D. et al. The lichen symbiosis re-viewed through the genomes of Cladonia grayi and its algal partner Asterochloris glomerata. BMC Genomics 20, 605 (2019).

80. Latorre, A. & Manzano-Marín, A. Dissecting genome reduction and trait loss in insect endosymbionts. Ann. N. Y. Acad. Sci. 1389, 52–75 (2017).

81. Vogel, K. J. & Moran, N. A. Functional and evolutionary analysis of the genome of an obligate fungal symbiont. Genome Biol. Evol. 5, 891–904 (2013).

82. González-Pech, R. A., Bhattacharya, D., Ragan, M. A. & Chan, C. X. Genome evolution of coral reef symbionts as intracellular residents. Trends Ecol. Evol. 34, 799–806 (2019).

83. van de Peppel, L. J. J. et al. Ancestral predisposition toward a domesticated lifestyle in the termite-cultivated fungus Termitomyces. Curr. Biol. 31, 4413–4421.e5 (2021).

84. Kajimura, H. & Hijii, N. Dynamics of the fungal symbionts in the gallery system and the mycangia of the ambrosia beetle,Xylosandrus mutilatus (Blandford) (Coleoptera: Scolytidae) in relation to its life history. Ecol. Res. 7, 107–117 (1992).

85. De Fine Licht, H. H. & Biedermann, P. H. W. Patterns of functional enzyme activity in fungus farming ambrosia beetles. Front. Zool. 9, 13 (2012).

86. Ibarra-Juarez, L. A., et al. Evidence for Succession and Putative Metabolic Roles of Fungi and Bacteria in the Farming Mutualism of the Ambrosia Beetle Xyleborus affinis. mSystems 5, (2020).

87. Hane, J. K., Williams, A. H., Taranto, A. P., Solomon, P. S. & Oliver, R. P. Repeat-induced point mutation: A fungal-specific, endogenous Mutagenesis process. in Fungal Biology 55–68 (Springer International Publishing, Cham, 2015). doi:10.1007/978-3-319-10503-1_4.

88. van Wyk, S., Wingfield, B. D., De Vos, L., van der Merwe, N. A. & Steenkamp, E. T. Genome-Wide Analyses of Repeat-Induced Point Mutations in the Ascomycota. Front. Microbiol. 11, 622368 (2020).

89. Teufel, F. et al. SignalP 6.0 predicts all five types of signal peptides using protein language models. Nat. Biotechnol. 40, 1023–1025 (2022).

90. Fischer, M. & Pleiss, J. The Lipase Engineering Database: a navigation and analysis tool for protein families. Nucleic Acids Res. 31, 319–321 (2003).

91. Rawlings, N. D. et al. The MEROPS database of proteolytic enzymes, their substrates and inhibitors in 2017 and a comparison with peptidases in the PANTHER database. Nucleic Acids Res. 46, D624–D632 (2018).

92. Buchfink, B., Reuter, K. & Drost, H.-G. Sensitive protein alignments at tree-of-life scale using DIAMOND. Nat. Methods 18, 366–368 (2021).

93. Zheng, J. et al. dbCAN3: automated carbohydrate-active enzyme and substrate annotation. Nucleic Acids Res. 51, W115–W121 (2023).

94. Pándy-Szekeres, G. et al. GPCRdb in 2023: state-specific structure models using AlphaFold2 and new ligand resources. Nucleic Acids Res. 51, D395–D402 (2023).

95. UniProt Consortium. UniProt: The universal protein knowledgebase in 2023. Nucleic Acids Res. 51, D523–D531 (2023).

96. Paradis, E. & Schliep, K. ape 5.0: an environment for modern phylogenetics and evolutionary analyses in R. Bioinformatics 35, 526–528 (2019).

97. Schliep, K. P. phangorn: phylogenetic analysis in R. Bioinformatics 27, 592–593 (2011).

98. van Wyk, S. et al. The RIPper, a web-based tool for genome-wide quantification of Repeat-Induced Point (RIP) mutations. PeerJ 7, e7447 (2019).

99. Mirarab, S. et al. ASTRAL: genome-scale coalescent-based species tree estimation. Bioinformatics 30, i541–8 (2014).

100. Kumar, S., Stecher, G., Li, M., Knyaz, C. & Tamura, K. MEGA X: Molecular Evolutionary Genetics Analysis across computing platforms. Mol. Biol. Evol. 35, 1547–1549 (2018).

101. Dos Reis, M. Dating microbial evolution with MCMCtree. Methods Mol. Biol. 2569, 3–22 (2022).

102. Tamura, K., Tao, Q. & Kumar, S. Theoretical foundation of the RelTime method for estimating divergence times from variable evolutionary rates. Mol. Biol. Evol. 35, 1770– 1782 (2018).

103. Kumar, S., Stecher, G., Suleski, M. & Hedges, S. B. TimeTree: A Resource for Timelines, Timetrees, and Divergence Times. Mol. Biol. Evol. 34, 1812–1819 (2017).

104. Bronson, A. W., Klymiuk, A. A., Stockey, R. A. & Tomescu, A. M. F. A Perithecial Sordariomycete (Ascomycota, Diaporthales) from the Lower Cretaceous of Vancouver Island, British Columbia, Canada. Int. J. Plant Sci. 174, 278–292 (2013).

105. Revell, L. J. phytools: an R package for phylogenetic comparative biology (and other things): phytools: R package. Methods Ecol. Evol. 3, 217–223 (2012).

